# Consolidation of reward memory during sleep does not require dopaminergic activation

**DOI:** 10.1101/703132

**Authors:** M Alizadeh Asfestani, V Brechtmann, J Santiago, J Born, GB Feld

## Abstract

Sleep enhances memories, especially, if they are related to future rewards. Although dopamine has been shown to be a key determinant during reward learning, the role of dopaminergic neurotransmission for amplifying reward-related memories during sleep remains unclear. In the present study, we scrutinize the idea that dopamine is needed for the preferential consolidation of rewarded information. We blocked dopaminergic neurotransmission, thereby aiming to wipe out preferential sleep-dependent consolidation of high over low rewarded memories during sleep. Following a double-blind, balanced, crossover design 20 young healthy men received the dopamine d2-like receptor blocker Sulpiride (800 mg) or placebo, after learning a Motivated Learning Task. The task required participants to memorize 80 highly and 80 lowly rewarded pictures. Half of them were presented for a short (750 ms) and a long duration (1500 ms), respectively, which enabled to dissociate effects of reward on sleep-associated consolidation from those of mere encoding depth. Retrieval was tested after a retention interval of 20 h that included 8 h of nocturnal sleep. As expected, at retrieval, highly rewarded memories were remembered better than lowly rewarded memories, under placebo. However, there was no evidence for an effect of blocking dopaminergic neurotransmission with Sulpiride during sleep on this differential retention of rewarded information. This result indicates that dopaminergic activation is not required for the preferential consolidation of reward-associated memory. Rather it appears that dopaminergic activation only tags such memories at encoding for intensified reprocessing during sleep.

## Introduction

Every day, the brain encodes large quantities of new information, and sleep related consolidation processes select the most relevant for long-term storage [1, 2]. During wakefulness, rewards play an important role to support this selection process, and, functional connectivity between the hippocampus and reward related areas at learning predicts memory retrieval a day later [3]. For this, the hippocampus which is initially involved in all episodic memory storage, and the reward centres, i.e., the ventral striatum and the ventral tegmental area (VTA) interact via a feedback loop [4] that enables dopamine to exert its influence on the learned behaviour [5]. However, while it seems clear that sleep plays an important role for the preferential consolidation of highly (over lowly) rewarded information [6–8], it remains open whether this effect depends on enhanced dopaminergic activation during sleep.

Sleep has been shown to support the consolidation of newly formed memories through the repeated replay of neuronal memory traces (e.g., [9–12]). It has been proposed that this replay also involves dopaminergic pathways, thereby, promoting better consolidation for the highly rewarded memories through enhanced neuroplasticity akin to processes acting during wakefulness [13]. This view is supported by findings in rats that underwent reward learning, where hippocampal replay was tightly linked to ventral striatal replay [14, 15]. Replay during sleep was also found in the VTA [16], thereby completing the hippocampal-ventral striatum-VTA loop implicated in this process. However, in another study, replay associated VTA activation remained restricted to post-encoding wakefulness and vanished during post-encoding sleep [17]. Thus, an alternative view assumes that rather than directly participating in sleep-dependent consolidation processes, dopamine activity elicited by rewards tags memory traces during encoding leading to more intense replay and accompanied plasticity during subsequent sleep. This view is supported by the finding in mice that optogenetically stimulating dopamine release in the hippocampus during encoding increases replay and consolidation of respective memories during subsequent sleep periods [18].

To collect causal evidence for or against a direct role of dopamine during sleep-dependent consolidation of reward-associated memories, we investigated, in humans, whether directly blocking dopamine interferes with the consolidation of such memories during sleep. In our Motivated Learning Task, sleep has been confirmed to preferentially consolidate memory for high rewarded pictures over low rewarded pictures [13]. Based on this evidence, here, we hypothesized that this difference would be wiped out, if dopaminergic transmission is blocked during sleep-dependent consolidation by administration of the dopamine d2-like receptor blocker Sulpiride.

## Methods

### Participants

Twenty healthy, native German speaking men fulfilling the requirements to enter higher education, aged on average 25.30 years (18-30 years) and with an average body mass index of 23.38 kg/m^2^ (20-25 kg/m^2^) completed this study. Details on the inclusion and exclusion criteria as well as general instructions can be found in the Supplementary Methods. Before the experimental nights, participants took part in an adaptation night under the same conditions of the experiment, which included the placement of the electrodes for polysomnographic recordings and of the cannula for the blood drawing. The ethics committee of the University of Tübingen approved the experiments. We obtained written informed consent from all participants before their participation.

### Design and procedures

The study followed a balanced, double-blind, placebo-controlled, within-subject crossover design (Figure 1A). Participants took part in two identical experimental sessions with the exception of administration of either Sulpiride (4 Dogmatil Forte, Sulpiride 200 mg, SanofiAventis, Germany) or placebo and parallel versions of the behavioural tasks where necessary, with at least two weeks interval between the sessions. The dose of 800 mg Sulpiride (p.o., plasma maximum: 3-6 h, plasma half-life: 6-10 h) was chosen, because at a lower dose Sulpiride is more likely to have an effect on presynaptic dopamine receptors and thus tends to increase dopamine release, whereas at 800 mg postsynaptic effects predominate. A single dose of 800 mg resulted in a 65% blockade of striatal d2-like receptors without adverse events in healthy volunteers [19]. Sulpiride was administered after the Learning Phase at 23:00, i.e., 15 min before lights off. We chose this timing in order to maximize drug levels during the slow wave sleep (SWS) rich first half of the night and thereby maximise the effects during occurrence of replay. The Retrieval Phase for the reward task was scheduled the next evening, i.e., as late as possible to minimize the residual amount of drug circulating at retrieval testing. An overview of the design can be found in Figure 1A and details are provided in the Supplementary Methods.

**Figure 1:**
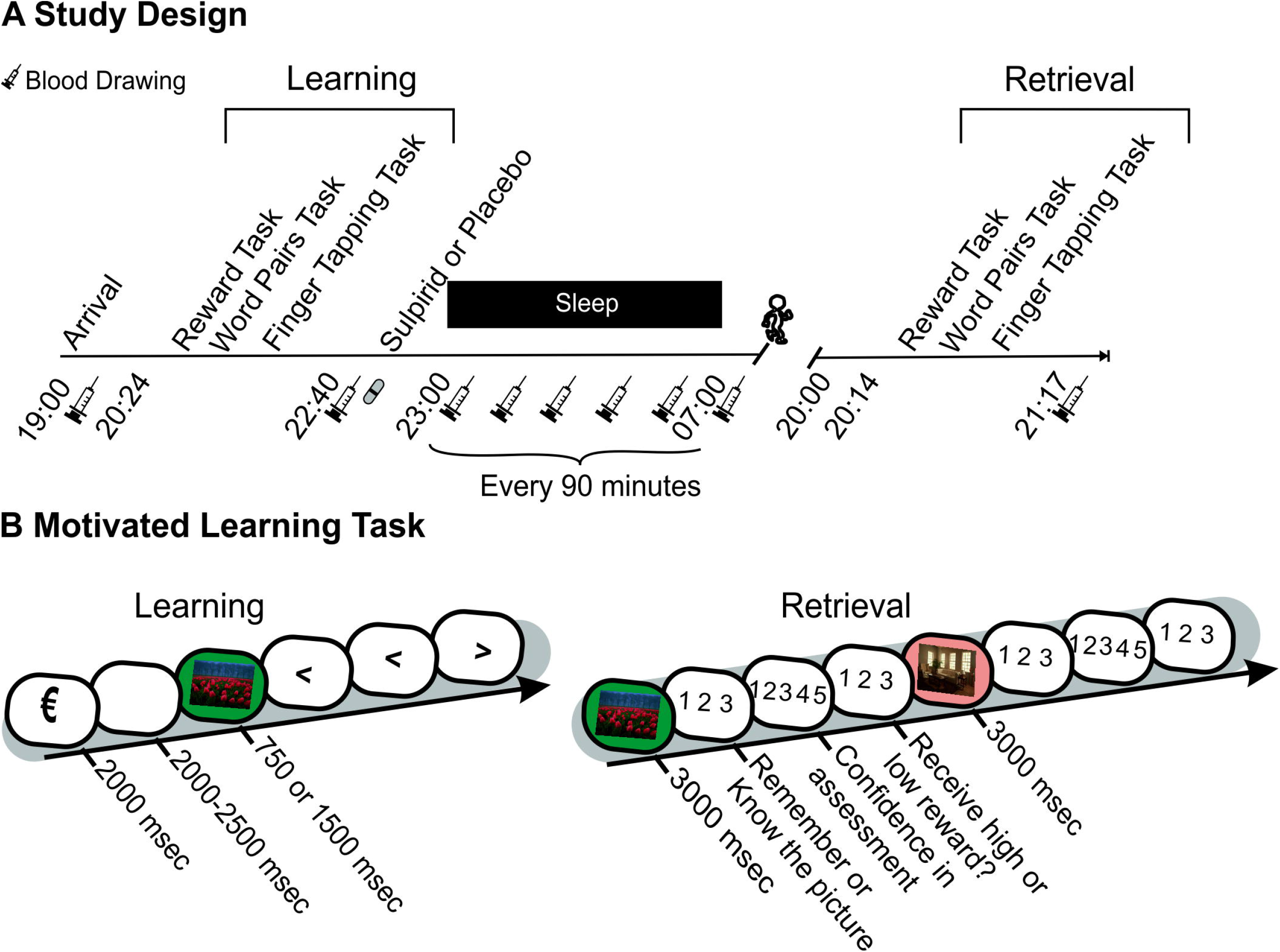
(A) Participants took part in two identical experimental sessions, but for the administration of placebo or Sulpiride. They started the session at around 7:00 pm, after preparing for blood sampling and sleep EEG, the Learning Phase started. Thereafter at 23:00, the capsules (Sulpirid or placebo) were orally administered. Participants were awakened at 7:15 the next morning. The retention interval was approximately 22 h, and retrieval was tested in the evening at approximately 20:00. Blood was drawn before and after learning, after retrieval, and in 1.5-h intervals during the night. (B) The motivated learning task was adapted from [3] and [13]. At learning participants were presented 160 pictures for 750 msec (short presentation) or 1500 msec (long presentation). Each picture was preceded by a slide indicating a high (1 Euro) or a low (2 Cents) reward for correctly identifying the picture at later recognition. After each picture, participants performed on three items of a distractor task, which afforded pressing the arrow key corresponding to the orientation of an arrow presented on the screen. At immediate (Learning Phase) and delayed recognition (Retrieval Phase) testing, participants were shown different sets of 80 new and 80 old pictures and had to identify them correctly, which earned them their reward (see Methods for details).

### Motivated Learning Task

The Motivated Learning Task was adapted from prior work [13] and required the participants to memorize 160 unique pictures of landscapes and living rooms in each of the two parallel versions (see Figure 1B for a visualisation and Supplementary Methods for details). Briefly, the pictures were preceded by a one Euro or a two Cents symbol to indicate high or low reward for successful later recognition. Pictures were also shown for a long (1500 msec) or a short (750 msec) duration to account for effects of encoding depth. During the two recognition sessions in the Learning and the Retrieval Phase subsets of the learned pictures were shown alongside sets of completely new pictures and participants had to recognize them. We used the sensitivity index d-prime as the primary outcome variable.

### Control measures

#### Behaviour

To control effects of the drug on declarative and procedural memory we used a word-pair task and a finger sequence tapping [20] task, respectively. In addition, we tested participants for long-term memory retrieval function (Regensburger Wortflüssigkeitstest [WFT]; [21]) during the Retrieval Phase. During the Learning and the Retrieval Phase we also measured vigilance [22], sleepiness [23] and mood [24]. Details about the control measures can be found in the Supplementary Methods.

#### Cortisol and prolactin

Serum concentrations of cortisol and prolactin were measured with the ADVIA Centaur XPT chemiluminescent immunoassay system from Siemens Healthineers, Eschborn, Germany. The inter-assay coefficients of variation were 5% for Cortisol and 2.5% for Prolactin. The area under curve was calculated as the weighted mean of the inter-interval approximation (time point n + time point (n + 1) / 2 × interval duration) for five time points, which occurred between lights out and waking, i.e., from 00:30 until 6:30.

### Polysomnography and sleep scoring

Polysomnography and sleep scoring were performed according to standard procedures [25]. Details can be found in the Supplementary Methods.

### Data reduction and statistical analysis

Three participants were excluded from the analysis, two of them for insufficient sleep and one for low levels of sleep and extremely long sleep latency. During blood sampling 73 draws (20.3 % of the total) were missed due to blockage of the tubing (occurring typically when the participant bends his arm during sleep). Singular missing values were replaced by interpolating between the neighbouring values. For two or more subsequent missing values, we calculated the average value of the rest of the participants at the same time point. Statistical analyses generally relied on ANOVAs (SPSS version 21.0.0 for Windows) including repeated-measures factors Treatment (Sulpiride vs. placebo), Reward (High vs. Low) and Duration (Long vs. Short). Of note, applying our previous analysis approach [13], we did not include a repeated measure factor for the Learning and Retrieval Phases as different stimuli were used for immediate and delayed recognition. Moreover, this would have led to a four factor ANOVA, which is hard to interpret. Significant interactions were followed up by post-hoc t-tests. Greenhous-Geisser correction of degrees of freedom was used, if data violated the assumption of homoscedacity.

## Results

### Motivated Learning Task

During the Learning Phase, highly rewarded pictures were recognized better than lowly rewarded pictures (main effect of reward: F_(1,16)_ = 25.03, p ≤ 0.001, Table 1 and Figure 2) and long duration pictures were recognized better than short duration pictures (main effect of duration: F_(1,16)_ = 6.75, p = 0.019). There were no significant interaction effects and no main effect of treatment in this analysis (all p > 0.511).

**Table 1:**
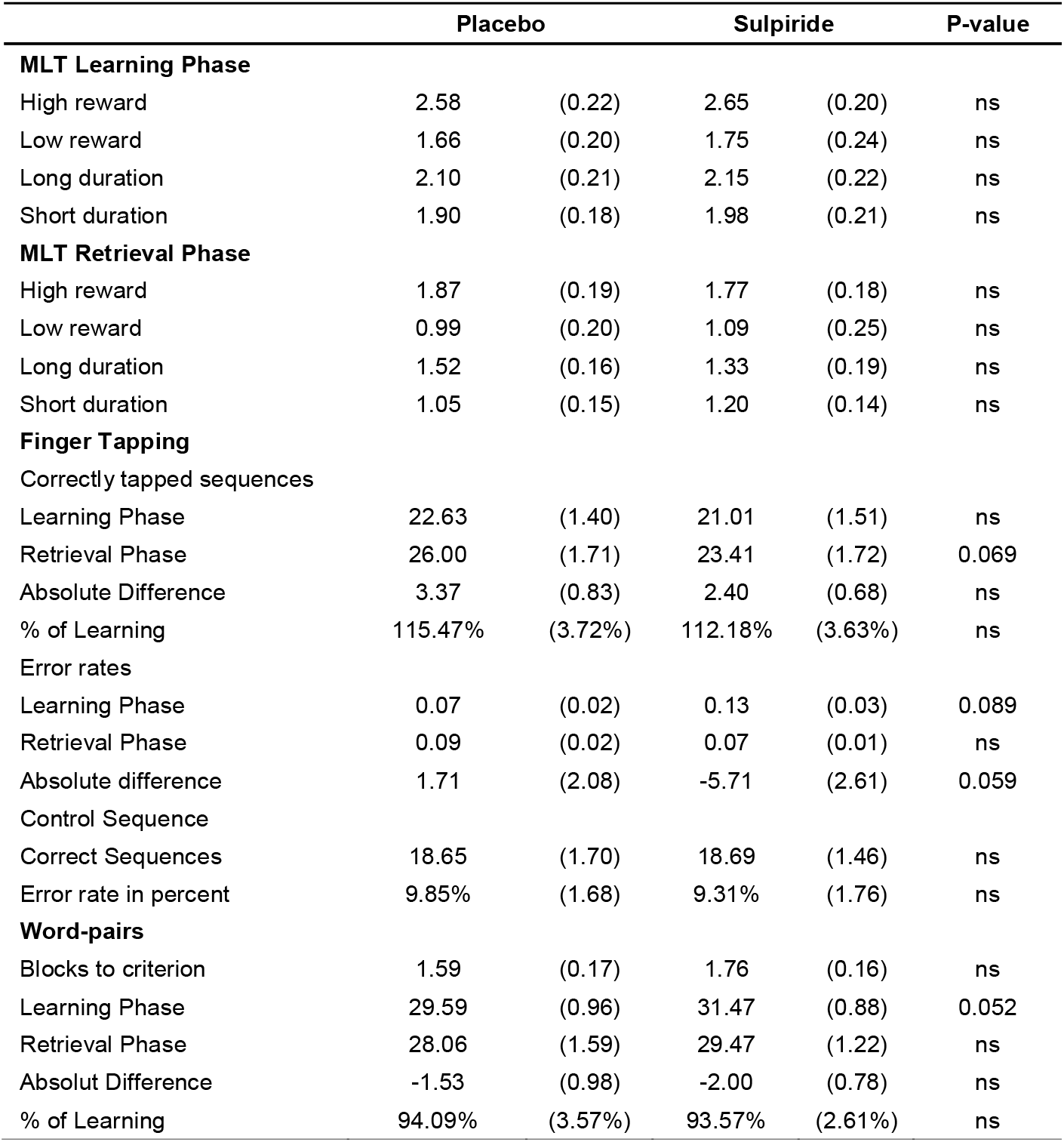
Memory Tasks. Mean (±SEM) values are provided for the Sulpiride and placebo conditions. Motivated Learning Task (MLT): d-prime is provided for performance during the Learning Phase and the Retrieval Phase. Finger tapping task: the average number of correctly tapped sequences per 30-sec trial and error rates (in percent of total tapped sequences) for finger sequence tapping during the last three 30-sec trials of the Learning Phase, the three trials during the Retrieval Phase and for the untrained control sequence. Additionally, the absolute difference (Retrieval-Learning) and percent of learning (Retrieval/Learningx100) are provided. Word-pair task: total amount of recalled words is given for the criterion trial during the Learning Phase and the recall trial during the Retrieval Phase. Also, the absolute difference (Retrieval-Learning) and percent of learning (Retrieval/Learningx100) are provided: **ns: p > .10.**

|  | Placebo |  | Sulpiride |  | P-value |
| --- | --- | --- | --- | --- | --- |
| MLT Learning Phase |  |  |  |  |  |
| High reward | 2.58 | (0.22) | 2.65 | (0.20) | ns |
| Low reward | 1.66 | (0.20) | 1.75 | (0.24) | ns |
| Long duration | 2.10 | (0.21) | 2.15 | (0.22) | ns |
| Short duration | 1.90 | (0.18) | 1.98 | (0.21) | ns |
| MLT Retrieval Phase |  |  |  |  |  |
| High reward | 1.87 | (0.19) | 1.77 | (0.18) | ns |
| Low reward | 0.99 | (0.20) | 1.09 | (0.25) | ns |
| Long duration | 1.52 | (0.16) | 1.33 | (0.19) | ns |
| Short duration | 1.05 | (0.15) | 1.20 | (0.14) | ns |
| Finger Tapping |  |  |  |  |  |
| Correctly tapped sequences |  |  |  |  |  |
| Learning Phase | 22.63 | (1.40) | 21.01 | (1.51) | ns |
| Retrieval Phase | 26.00 | (1.71) | 23.41 | (1.72) | 0.069 |
| Absolute Difference | 3.37 | (0.83) | 2.40 | (0.68) | ns |
| % of Learning | 115.47% | (3.72%) | 112.18% | (3.63%) | ns |
| Error rates |  |  |  |  |  |
| Learning Phase | 0.07 | (0.02) | 0.13 | (0.03) | 0.089 |
| Retrieval Phase | 0.09 | (0.02) | 0.07 | (0.01) | ns |
| Absolute difference | 1.71 | (2.08) | -5.71 | (2.61) | 0.059 |
| Control Sequence |  |  |  |  |  |
| Correct Sequences | 18.65 | (1.70) | 18.69 | (1.46) | ns |
| Error rate in percent | 9.85% | (1.68) | 9.31% | (1.76) | ns |
| Word-pairs |  |  |  |  |  |
| Blocks to criterion | 1.59 | (0.17) | 1.76 | (0.16) | ns |
| Learning Phase | 29.59 | (0.96) | 31.47 | (0.88) | 0.052 |
| Retrieval Phase | 28.06 | (1.59) | 29.47 | (1.22) | ns |
| Absolut Difference | -1.53 | (0.98) | -2.00 | (0.78) | ns |
| % of Learning | 94.09% | (3.57%) | 93.57% | (2.61%) | ns |

**Figure 2:**
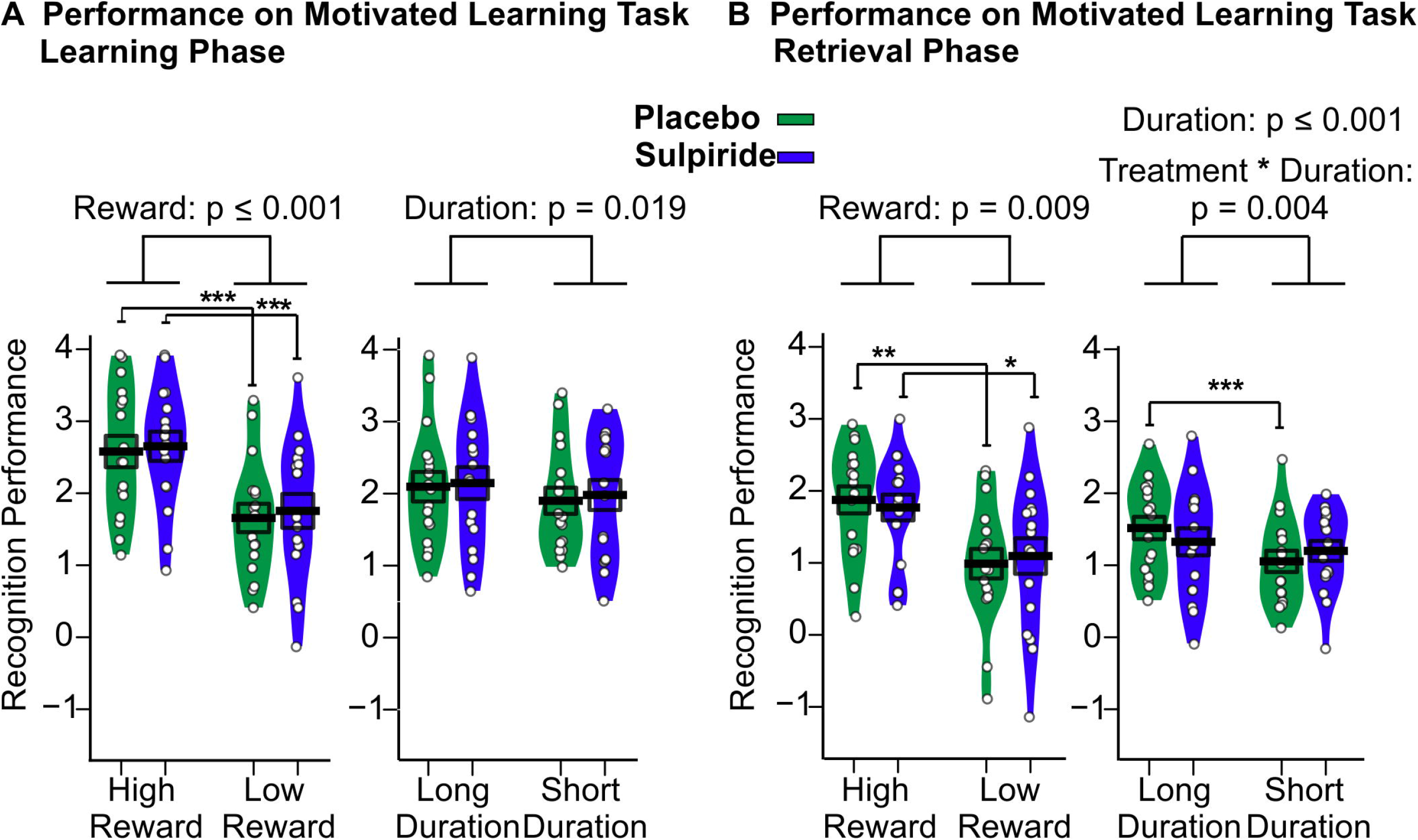
**(A)** Performance on the motivated learning task for the immediate recognition test during the Learning Phase before sleep and **(B)** delayed recognition test during the Retrieval Phase after sleep for the Sulpiride (purple) and the placebo (green) conditions. Mean (±SEM) performance is indicated as d-prime, that is, the z value of the hit rate minus the z value of the false alarm rate. n = 17. **\*\*\*p≤ .001, **p≤ .01 and *p≤ .05**

During the Retrieval Phase, highly rewarded and longer duration pictures were recognized significantly better than lowly rewarded and short duration pictures, respectively (main effect of reward: F_(1,16)_ = 8.94, P = 0.009; main effect of duration: F_(1,16)_ = 20.54, p ≤ 0.001). However, there was no evidence of Sulpiride affecting the recognition performance in general (main effect of treatment: F _(1,16)_ = 0.02, p =0.892) or recognition performance in the reward conditions differentially (Treatment × Reward: (F_(1,16)_ = 0.59, p =0.454). To test the robustness of this null effect an exploratory overall analysis including Learning Phase and Retrieval Phase data was conducted, which also did not yield an effect of Sulpiride regarding high or low rewards (Treatment × Reward: F_(1,32)_ =0.57, p = 0.460). Rather we found that Sulpiride diminished the performance difference between long and short duration pictures during the Retrieval Phase (Medication × Duration: F_(1,16)_ =11.06, p = 0.004). In the placebo condition long duration items were recognized better than short duration items (Long Duration: mean = 1.52, SD = 0.64, Short Duration: mean = 1.05, SD = 0.61, t_(16)_ = 6.23, p ≤ 0.001), which was not true for the Sulpiride condition (Long Duration: mean = 1.33, SD = 0.77, Short Duration: mean = 1.20, SD = 0.57, t_(16)_ = 1.44, p =0.170, Figure 2).

To determine response strategies we calculated the response bias, i.e., the negative mean of the z-value of the hit- and of the false-alarm rate. In both recognition phases participants’ reactions were more conservative for the high-reward pictures (Learning Phase: F_(1,16)_ = 20.02, p ≤ 0.001, Retrieval Phase: F_(1,16)_ = 6.68, p = 0.020, Table 2). None of the other contrasts were significant (all p > 0.193).

**Table 2:** Motivated Learning Task Additional Response Information. Mean (±SEM) values are given for the Sulpiride and placebo conditions for hits, false alarms and response bias during the Learning Phase and the Retrieval Phase. ns: p > .10.

|  | Placebo |  | Sulpiride |  | P-value |
| --- | --- | --- | --- | --- | --- |
| Hits |  |  |  |  |  |
| Learning Phase |  |  |  |  |  |
| High reward | 0.79 | (0.04) | 0.82 | (0.03) | ns |
| Low reward | 0.76 | (0.04) | 0.79 | (0.03) | ns |
| Long duration | 0.80 | (0.03) | 0.82 | (0.03) | ns |
| Short duration | 0.75 | (0.04) | 0.78 | (0.03) | ns |
| Retrieval Phase |  |  |  |  |  |
| High reward | 0.69 | (0.05) | 0.63 | (0.04) | ns |
| Low reward | 0.64 | (0.04) | 0.64 | (0.04) | ns |
| Long duration | 0.73 | (0.04) | 0.65 | (0.05) | 0.028 |
| Short duration | 0.59 | (0.05) | 0.62 | (0.04) | ns |
| False Alarms |  |  |  |  |  |
| Learning Phase |  |  |  |  |  |
| High reward | 0.07 | (0.02) | 0.08 | (0.03) | ns |
| Low reward | 0.23 | (0.05) | 0.25 | (0.05) | ns |
| Retrieval Phase |  |  |  |  |  |
| High reward | 0.14 | (0.03) | 0.12 | (0.03) | ns |
| Low reward | 0.31 | (0.06) | 0.29 | (0.06) | ns |
| Response Bias |  |  |  |  |  |
| Learning Phase |  |  |  |  |  |
| High reward | 0.40 | (0.08) | 0.30 | (0.10) | ns |
| Low reward | 0.02 | (0.11) | -0.01 | (0.11) | ns |
| Retrieval Phase |  |  |  |  |  |
| High reward | 0.39 | (0.14) | 0.52 | (0.11) | ns |
| Low reward | 0.09 | (0.13) | 0.15 | (0.11) | ns |

We also separately analysed hit rates and false alarm rate, which largely paralleled data for the sensitivity index. Hit rates were higher for longer duration pictures (Learning Phase: F_(1,16)_ = 7.48, p = 0.015, Retrieval Phase: F_(1,16)_ = 21.83, p ≤ 0.001) and highly rewarded pictures (Learning Phase: F_(1,16)_= 6.94, p = 0.018, Retrieval Phase: F _(1,16)_ = 6.40, p = 0.022) and at learning false alarms were reduced for highly rewarded pictures (Learning Phase: F_(1,16)_ =12.83, p = 0.002). For the hit rate, we also found that Sulpiride differentially affected performance for long and short duration items, during the Retrieval Phase (Treatment × Duration: F_(1,16)_ = 11.86, p =0.003; see Table 2).

### Control Measures

No systematic evidence was found for the drug affecting declarative and procedural memory tasks (Table 1 for descriptive data and Supplementary Results for details). The other control measures also did not provide evidence for large systematic effects of Sulpiride (Supplementary Results for details).

#### Cortisol and prolactin

We found no evidence of Sulpiride affecting cortisol levels in general (Treatment: F =1.65, p = 0.22) or at specific time points (Treatment x Time point: F_(1,16)_ = 0.92, p = 0.45). However, prolactin levels were increased in the Sulpiride condition at some time points (Treatment: F _(1,16)_ = 227.00, p ≤ 0.001, Time point: F_(1,16)_ = 40.49, p ≤ 0.001, Treatment × Time point: F_(1,16)_ = 37.32, p ≤ 0.001). This was true for all samples from 00:30 until 21:30 (post-hoc t-test all p ≤ 0.001) as well as in an analysis of the area under the curve (AUC) from 00:30 until 06:30 (t_(16)_ = −16.81, p ≤ 0.001, see Figure 3 C). This effect can be explained by Dopamine having a strong inhibitory effects on prolactin secretion [26]. Since prolactin was still elevated during the Retrieval Phase in the Sulpiride condition, it is likely that an active level of the drug remained. This may explain lowered reaction speed and positive mood in the Sulpiride group at this time point. Since our timing of the Retrieval Phase was already maximally postponed after intake this residual amount of drug cannot be prevented in our paradigm.

**Figure 3:**
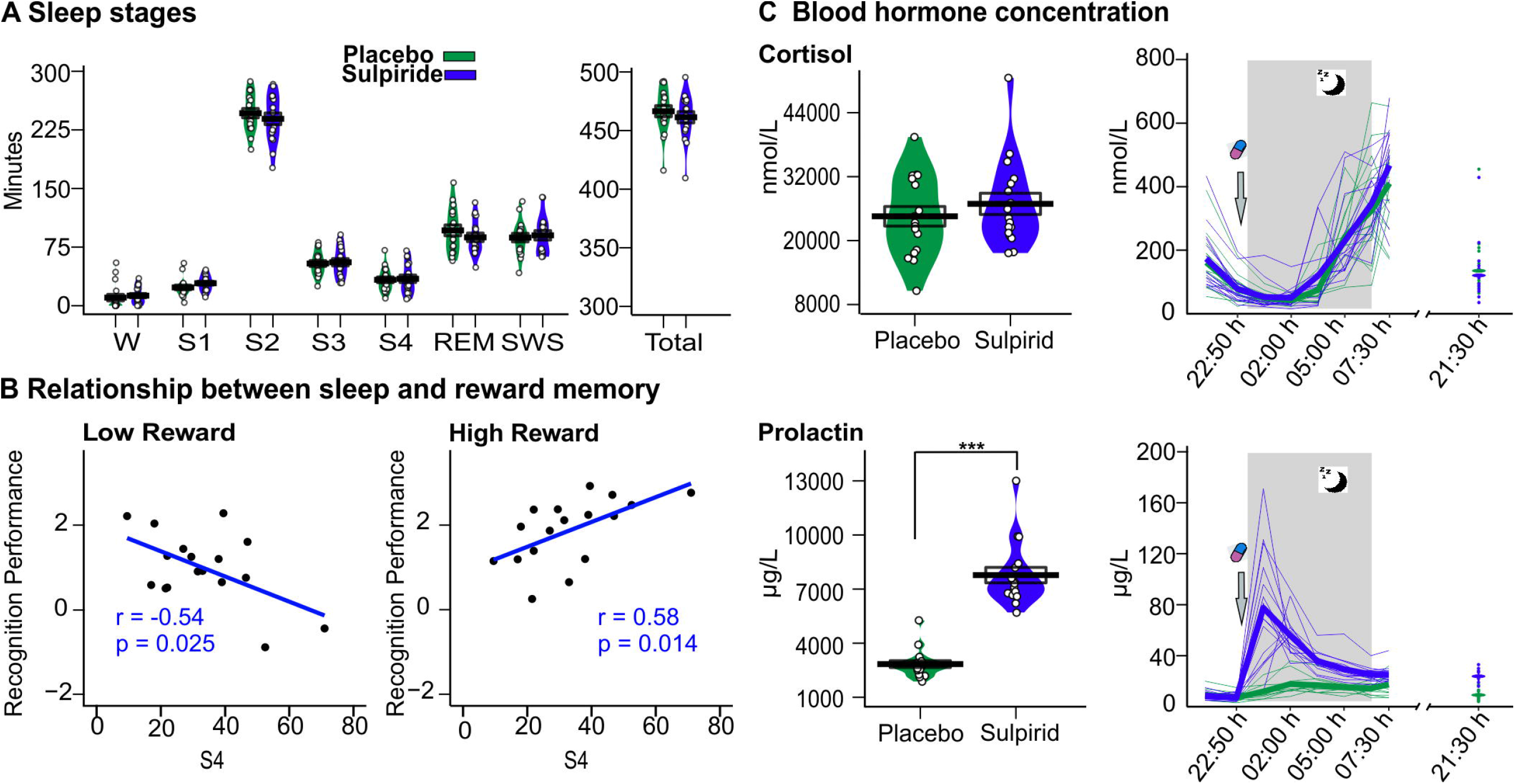
**(A)** Sleep stages. Mean (±SEM) time (in minutes) spent in non-REM Sleep Stages S1, S2, S3, and S4; in REM sleep; in SWS (i.e., the sum of S3 and S4) and total sleep time, are provided for the Sulpiride (purple) and placebo (green) conditions. **(B)** Correlation, across placebo condition between sleep stage 4 and low as well as high rewarded memories, respectively. **(C)** Blood hormone concentration. Values for Cortisol and Prolactin are shown at the top and bottom, respectively. Mean (±SEM) area under the curve (AUC) (from 00:30 until 06:30) is shown on the left and mean (thick lines) and individual data (thin lines) per time point is shown on the right. The Sulpiride condition is shown in purple and placebo is shown in green. **\*\*\*p≤ .001**

### Sleep Parameters

Total sleep time and time spent in the different sleep stages did not significantly differ between the treatment conditions (all p ≥ 0.199, see figure 3A). In post-hoc analyses, we explored correlations between sleep parameters and performance on the reward memory task in the placebo condition (Figure 3B). We found a significant positive correlation between the time spent in sleep stage 4 and Retrieval Phase recognition performance for highly rewarded pictures (r = 0.58, p = 0.014), whereas this relationship was negative for lowly reward pictures (r = −0.54, p = 0.025). Meaning that participants generally performed better on highly rewarded picture recognition and worse on lowly rewarded picture recognition the more sleep stage 4 they had. This relationship remained largely consistent but was slightly weaker, when data for both conditions were pooled with similar correlations between the time spent in sleep stage 4 and Retrieval Phase performance (highly rewarded pictures: r =0.50, p = 0.041, lowly rewarded pictures: r = −0.48, p = 0.053).

## Discussion

We investigated whether activation of the dopaminergic reward network during sleep is necessary for selective consolidation of highly over lowly rewarded memories. To this end, we blocked dopamine d2-like-receptors – using the selective antagonist Sulpiride – during sleep after participants learned a set of highly or lowly rewarded pictures. We found that, generally, highly rewarded pictures were retained better than lowly rewarded pictures across sleep, which concurs with earlier reports [3, 13] and is also in line with reports of sleep preferentially enhancing the retention of highly over lowly rewarded information (e.g., [7]). Contrary to our hypothesis, Sulpiride did not affect these reward related differences in retention. Rather, we found that Sulpiride diminished the preferential retention of deeply over shallowly encoded pictures. Importantly, the dopaminergic receptor antagonist did not significantly alter sleep architecture. Together, these findings exclude a causal contribution of dopaminergic activation during sleep to the preferential consolidation of reward-associated memory.

Both in the Sulpiride and the placebo condition, participants recognized highly rewarded pictures better than lowly rewarded pictures at retrieval testing after sleep. With respect to previous studies, this finding reflects the successful involvement of midbrain dopaminergic structures during the encoding of reward related information in the hippocampus by our Motivated Learning Task [27–29], which is eventually necessary for sleep to selectively enhance highly rewarded information [6–8]. There is overwhelming evidence that this sleep-dependent consolidation relies on the replay of neuronal memory traces during slow wave sleep (e.g., [9, 10, 30, 31]). In addition, some studies suggested that the reward circuitry of the brain, i.e., the hippocampus-ventral striatum-ventral tegmental area-hippocampus loop, participates in this replay [14–16,32]. However, as our data revealed, a potent block of dopaminergic neurotransmission using Sulpiride does not block the enhanced consolidation of highly over lowly rewarded information and, thus, the dopaminergic reward circuits seem not to engage in this consolidation process. This finding agrees with a recent study of single unit recordings in the hippocampus and VTA of rats, which learned reward locations in a maze [17]. Here, replay during quiet wakefulness directly after task performance showed a co-involvement of hippocampus and VTA, whereas this relation was not evident for replay during subsequent slow wave sleep. Another study in rats showed that dopaminergic activation during learning can enhance replay during sleep even in the absence of a behavioural effect at learning [18]. Those findings in combination with the present data, support the idea that augmented neuronal replay, rather than co-activation of dopaminergic neurotransmission, is the mayor player enhancing memory consolidation for highly rewarded information during sleep.

At a first glance, the present results are at odds with our study where the dopamine d2-like receptor agonist Pramipexole selectively enhanced sleep-dependent consolidation of lowly rewarded pictures in the same task [13]. However, unlike Sulpiride, Pramipexole administration caused severe disturbances of sleep. In fact, in mice, optogentically activating dopaminergic neurons of the VTA was found to promote wakefulness, whereas inhibition of the same cells supressed wakefulness, even in the presence of highly appetitive or threatening stimuli [33]. Against this backdrop, it seems prudent to interpret the effects of Pramipexole in that study as non-physiological, i.e., assuming that the enhancing effect the drug had on low reward items was secondary to its arousing effects.

Our additional post-hoc correlation analyses of the placebo condition revealed further hints consistent with a role of replay in specifically enhancing highly rewarded information. Here, time spent in deepest slow wave sleep (i.e., sleep stage 4) positively correlated with recognition performance of highly rewarded pictures, but negatively with performance on low-reward pictures. Replay has been especially connected to consolidation during slow wave sleep and sleep stage 4 has the most slow oscillations (of all sleep stages). These are thought to drive spindles top-down and, eventually memory replay activity together with ripples in hippocampal networks [12, 34, 35]. Ripples together with spindles are likely the oscillations, which promote the neuroplasticity that strengthens memory traces in this process [36–39]. Importantly, hippocampal ripples appear to be simultaneously involved in processes of synaptic downscaling and forgetting [1, 40, 41], and, thus, represent a putative mechanism explaining our observation that time in stage 4 sleep was also negatively correlated with recognition of low-reward items.

Our finding that the enhanced recognition of highly rewarded pictures was already strongly evident at immediate recall, during the Learning Phase, points towards the dopaminergic system exerting its enhancing role on rewarded information already during learning [29, 42]. Although some studies suggests that rewards mainly enhance memory performance after a delay rather than directly [13, 43–45]. What is important here is that this reward effect during the Learning Phase cannot be taken as evidence that preferential consolidation of highly rewarded information occurs in relation to encoding strength alone, as our task also included pictures that were shown for a short or a long duration. This also led to a recognition advantage for long duration pictures during the Retrieval Phase that, however, was wiped out by Sulpiride during sleep. This finding opens the possibility that dopamine plays a non-reward related role during sleep, possibly in relation to recently discovered post-encoding memory enhancement of novel stimuli by release of dopamine in the hippocampus that is mediated by the locus coeruleus [46], a brain region with activity regulated by the sleep slow oscillation [47]. Of note, this finding was not predicted before conducting our study and Takeuchi and colleagues tested blocking d1-like rather than d2-like receptors in the hippocampus. So future research will have to scrutinize these effects.

A limitation of our study is that we blocked d2-like dopamine receptors and therefore finding no interaction between treatment and reward consolidation does not rule out that d1-like receptors play a more important role during sleep. Considering evidence that both d2-like and d1-like receptors are implicated in hippocampus dependent tasks and reward learning [48–51] future studies should focus on d1-like receptor related effects using drugs like L-dopa, or dietary dopamine depletion [52] during sleep-dependent consolidation.

In conclusion, our data challenge the idea that replay during sleep engages dopaminergic inputs to the hippocampus via a feedback loop consisting of the brain’s reward centres to selectively enhance information related to high rewards. Rather, it seems likely that a form of dopamine related tagging occurs at encoding that enhances replay activity for relevant memories during sleep, thereby, strengthening them.

## Supporting information

Supplementary Methods and results

## Acknowledgements

The authors would like to thank Joachim Ingenschay for assisting data collection.

