## Supplementary Methods and results for "Consolidation of reward memory during sleep does not require dopaminergic activation"

**Participants**

Before entering the study, all participants underwent a routine medical examination to exclude any psychiatric, neurological, cardiovascular, endocrine or gastrointestinal diseases. Participants with hypersensitivity to Sulpiride or Benzamide derivatives, regular excessive alcohol consumption (regularly more than two bottles of beer per day), nicotine consumption or taking regular medication (i.e., including painkillers and sleeping pills) were excluded. The medical screening relied on a structured interview asking for current or past diagnosed conditions, a physical examination as well as a blood pressure and a routine blood screening test (including haemoglobin, sodium, potassium, calcium, chloride, glucose, bilirubin, glutamate pyruvate transaminase, alkaline phosphatase, Gamma-glutamyl-trans-peptidase, C-reactive protein, Partial thromboplastin time) and only healthy participants were included. In addition, participants reported a normal sleep-wake cycle, no shift work, night work or intercontinental flights (>4 h time difference) for at least 6 weeks before the experiments. Participants were instructed to keep a regular sleep schedule in the week before the experiment (approximately sleeping from 23:00 to 7:00 each night), to go to bed at 23:00 the night before experiments, to get up at 07:00 on experimental days and, during these days, not to take any naps, not to drink caffeine-containing drinks after 13:00 and also not to consume alcohol starting one day before the experimental nights. Adherence to these rules was assessed with a questionnaire at the very beginning on each experimental session.

**Design and procedures**

On the experimental nights, participants arrived at 19:00 filled a general questionnaire and then an intravenous cannula was placed for drawing blood. Afterwards, electrodes were applied for polysomnographic recordings. Next, they filled in the questionnaires on mood and sleepiness. About one hour after cannula placement, the first blood sample was taken and then the behavioural tasks were performed. First they performed a vigilance task and then the reward task was learned with immediate recall of half the items scheduled after a 15 min break. Then, after additional breaks of 5 min each, declarative and procedural contents were learned as control tasks. Afterwards they again preformed the vigilance task and filled in the questionnaires on mood and sleepiness. At 22:50, blood was sampled again and at 23:00 the participant orally received either Sulpiride or placebo. Participants slept from 23:15 to 7:15 and a polysomnogram was recorded. During the night, blood was sampled every 1.5 h starting at 0:30. A long thin tube connected to the cannula enabled blood collection during the night from an adjacent room without disturbing the participant’s sleep. Participants were woken between 7:00 and 7:30 preferably from sleep stages 1 or 2. Next, they filled in mood questionnaires. Blood was sampled again approximately 15 min after waking up. Then participants were allowed to shower and received a standardized breakfast (2 slices of bread, butter, cheese and water) before leaving the laboratory. In the evening of the same day, participants returned to the laboratory at 20:00 and filled in the mood and sleepiness questionnaires. Afterwards, they performed the vigilance task and then the word fluency task. Next they performed the finger sequence tapping task (first retrieval of the learned sequence was tested and then a new control sequence was learned). After 5 min of break each, they were asked to retrieve the declarative word-pair task and to recognize the reward contents, respectively. Then they performed the vigilance task and answered the mood and sleepiness questionnaires again. Blood was sampled once more at 21:30 before participants left the lab.

**Motivated Learning Task**

Presentation of 80 highly rewarded pictures was preceded (delay 2000 to 2500 msec) by a 1 euro symbol whereas the other 80 lowly rewarded pictures were preceded by a 2 cents symbol, and participants were informed that they would receive the respective reward for every hit during subsequent recognition. They were also informed that a correct rejection (identifying a novel picture as not being presented at learning) at recognition testing would earn them 51 cents and that for a miss or a false alarm they would lose 51 cents. This was done to exclude potential strategy effects, for example, only choosing items that would yield high rewards as old. Forty pictures each of the two reward conditions were presented for 750 and 1500 msec, respectively, to control for effects of encoding depth. Encoding depth was manipulated as the reward manipulations may also lead to differences in encoding depth, and we were interested whether the effect of Sulpiride would be independent of this confound. Each picture was followed by three items of a distraction task where participants had to press one of two buttons according to the orientation of an arrow presented on the screen, and 1 sec later, the next trial started. Participants were allowed to train the task for three items including the recognition procedure before learning the pictures, and the first two and last two pictures that were added in addition to the 160 pictures were excluded from later recognition testing to buffer recency and primacy effects. Participants were also informed that recognition would be tested twice, immediately after learning and in the evening of the next day. Immediate recognition started 15 min after learning had finished and, before starting, participants were reminded of the reward contingencies (also by training on three pictures). During recognition testing they were shown 80 of the original pictures together with 80 new pictures in a pseudo-random order and asked to indicate for each picture if they remembered or knew the picture (correct answers were summed and used to calculate individual hit rates) or if it was new by pressing a key on the keyboard (1, 2, or 3, respectively). They also pressed a key (1 or 2, respectively) according to whether they believed to receive a high or a low reward for the answer (thus, incorrect remember and know judgments allowed us to calculate individual false alarm rates for high and low reward categories). All participants received mock feedback of how much they had earned after each recognition test (the message “You performed slightly above average and will receive X euros” was displayed with amounts varying between 47.5 and 52.5 euros”). This was done to keep participants motivated while controlling effects of high or low performance. Delayed recognition that was performed the next evening was identical, but the other 80 learned pictures were used and 80 completely new pictures were shown in comparison. D-prime, that is, the z-value of the hit rate minus the z-value of the false alarm rate was calculated as dependent variable, which is independent of response strategies. For constructing task stimuli, 32 similar groups of 20 pictures each were generated with regard to mean valence and arousal ratings as assessed in a pilot study (n = 5). The presentation of the groups was then balanced across the old/new, immediate/delayed recognition, short/long presentation, and high/low reward conditions for the different participants.

**Control measures**

In the declarative control task participants learned a list of 40 associated word-pairs (e.g., Painter - Pianist, presented for 4 sec each). After viewing the complete list of 40 pairs in a random order participant’s performance was tested using a cued recall procedure. After each response, the complete pair was displayed for 2 sec. This procedure was repeated until the participant reached 60% correct responses. The same cued recall procedure was used once more during the Retrieval Phase, except that no feedback of the correct answer was given. To measure the overnight retention we calculated the absolute differences between word-pairs recalled during the Retrieval Phase and word-pairs recalled during the last run of the Learning Phase. For the procedural finger sequence tapping task participants had to repeatedly input a 5-element sequence (e.g., 4-1-3-2-4 or 4-2-3-1-4) with the fingers of their non-dominant hand as fast and as accurately as possible. This had to be done during twelve 30-sec trials interrupted by 30-sec breaks. We scored for speed (number of correctly completed sequences) and error rate (proportion of incorrectly tapped sequences). Learning performance was calculated by averaging performance for the last three of these trials. During the Retrieval Phase, participants performed another three trials, which were also averaged. The absolute differences between the Retrieval Phase and performance in the Learning Phase was calculated as a measure of overnight retention. As a control for effects of the drug during the Retrieval phase, participants performed the trials of a novel unlearned control sequence.

During the Retrieval Phase, participants were also tested on a word generation task to control for effects of the drug on long-term memory retrieval function. Within 2 min each, participants had to generate as many words as possible first starting with a specified letter (p or m) and then from a specified category (jobs or hobbies). Further control measures were tested before and after the Learning Phase, as well as, the Retrieval Phase. We measured participants’ vigilance with a 5-min version of the psychomotor vigilance task (PVT), their mood with the Positive and Negative Affective Schedule (PANAS) and their subjective sleepiness with the Stanford Sleepiness Scale (SSS). After finishing each session, participants were asked if they believed to have received Sulpiride or placebo.

**Polysomnography and sleep scoring**

The EEG was recorded continuously from electrodes (Ag-AgCl) placed according to the 10–20 System, referenced to two linked electrodes attached to the mastoids. EEG signals were filtered between 0.16 and 35 Hz and sampled at a rate of 250 Hz using a Brain Amp DC (Brain Products GmbH, Munich, Germany). Additionally, horizontal and vertical eye movements (HEOG, VEOG) and the EMG (via electrodes attached to the chin) were recorded for standard polysomnography. Sleep architecture was determined according to standard polysomnographic criteria using EEG recordings from C3 and C4 (Rechtschaffen & Kales, 1968). Scoring was carried out independently by two experienced technicians who were blind to the assigned treatment. Differences in scoring between the scorers were resolved by consulting a third experienced technician. For each night, total sleep time and time spent in the different sleep stages (wake; Sleep Stages 1, 2, 3, 4; SWS, that is, sum of Sleep Stages 3 and 4; REM sleep) was calculated in minutes.

**Supplementary Results**

***Declarative and procedural memory tasks.*** In the declarative word-pair task we found no effect of Sulpiride on retention (t _(1,16)_= 0.35, p = 0.729). Under Sulpiride, participants recalled significantly less word-pairs during the Retrieval Phase than during the Learning Phase (t_(16)_=2.56, p = 0.021). However, this difference was already apparent during the Learning Phase (t_(16)_= -2.10, p = 0.052, Table 1). There was no difference between the treatments regarding the amount of runs needed to reach the learning criterion (t_(16)_= -1.38, p = 0.188).

In the finger tapping task, there was a trend wise effect for error rates decreasing more in the Sulpiride condition across the retention interval (t_(16)_ = 2.03, p = 0.059). However, this was from a trend wise higher baseline in the Learning Phase (t_(16)_= -1.81, p = 0.089). For the correctly tapped sequences we found a trend wise effect for participants tapping less correct sequences in the Sulpiride condition during the Retrieval Phase (t_(16)_ = 1.95, p = 0.069). There was no effect of Sulpiride on the control sequence only performed during the Retrieval Phase (correct sequences: t _(16)_= - 0.02, p = 0.982; error rates: t_(16)_ = 0.25, p = 0.809).

***Word fluency, vigilance, mood, and subjective sleepiness.*** Descriptive data can be found in Supplementary Table 1. We did not find any significant differences in long-term memory retrieval performance (as measured by the word fluency task, all p≥ 0.868). In the vigilance task (PVT) we found significantly higher reaction speed (i.e., reaction time-1) in the Placebo condition after the Retrieval Phase (t_(16)_ = 3.13, p = 0.006, all other p ≥ 0.637). In the placebo condition compared to the Sulpiride condition, the mood questionnaire (PANAS) showed significantly higher positive mood before the Retrieval phase (t_(16)_= 2.25, p = 0.039) and a trend toward more negative mood after the Learning Phase (t_(16)_= 2.06, p = 0.056). In the Sulpiride condition compared to the placebo condition, there was some evidence for increased subjective sleepiness (SSS) after the Learning Phase (t_(16)_ = -2.75, p = 0.014) and a trend toward increased sleepiness before the Retrieval Phase (t_(16)_ = -1.81, p = 0.090). Participants were not able to discriminate between Sulpiride and placebo (McNemars’ exact test: p ≥ 0.791).

**Supplementary Table 1: Control Measures.** Mean (±SEM) values are provided for the Sulpiride and placebo conditions. SSS = Stanford Sleepiness Scale (subjective sleepiness); PANAS = Positive and Negative Affective Scale (mood); PVT = Psychomotor Vigilance Task (reaction speed = 1/[RT in msec]); WFT = Word Fluency Test (Regensburger Wortfluessigkeitstest) long-term retrieval capabilities**. ns: p > .10.**

|  | Placebo | | Sulpirid | | P-value |
| --- | --- | --- | --- | --- | --- |
| SSS |  |  |  |  |  |
| Before Learning | 2.94 | (0.16) | 2.94 | (0.26) | ns |
| After Learning | 3.71 | (0.27) | 4.47 | (0.26) | 0.014 |
| Before retrieval | 2.29 | (0.22) | 3.00 | (0.32) | 0.090 |
| After Retrieval | 2.71 | (0.19) | 3.00 | (0.33) | ns |
| Positive Affect (PANAS) |  |  |  |  |  |
| Before Learning | 2.42 | (0.09) | 2.48 | (0.13) | ns |
| After Learning | 1.91 | (0.15) | 1.77 | (0.13) | ns |
| Before retrieval | 2.58 | (0.16) | 2.18 | (0.15) | 0.039 |
| After Retrieval | 2.32 | (0.12) | 2.15 | (0.14) | ns |
| Negative Affect (PANAS) |  |  |  |  |  |
| Before Learning | 1.08 | (0.03) | 1.06 | (0.04) | ns |
| After Learning | 1.04 | (0.01) | 1.01 | (0.01) | 0.056 |
| Before retrieval | 1.02 | (0.01) | 1.01 | (0.01) | ns |
| After Retrieval | 1.02 | (0.03) | 1.01 | (0.01) | ns |
| PVT |  |  |  |  |  |
| Before Learning | 3.56 | (0.09) | 3.59 | (0.07) | ns |
| After Learning | 3.45 | (0.10) | 3.45 | (0.09) | ns |
| Before retrieval | 3.54 | (0.10) | 3.55 | (0.08) | ns |
| After Retrieval | 3.53 | (0.10) | 3.33 | (0.07) | 0.006 |
| WFT |  |  |  |  |  |
| Category | 20.18 | (0.91) | 20.24 | (1.26) | ns |
| Letter | 19.94 | (1.01) | 20.12 | (1.39) | ns |
